# Camera trap monitoring of unmarked animals: a map of the relationships between population size estimators

**DOI:** 10.1101/2025.02.28.640755

**Authors:** Clément Calenge

## Abstract

The use of camera traps to monitor unmarked animal populations has expanded during the last decade, leading to the development of several density estimation methods. This plethora of methods may be confusing for the newcomer to the field. Some methods, such as the random encounter model, require the knowledge of the mean travel speed of the animals, while others, such as camera trap distance sampling, do not rely on such assumptions. Different methods, like instantaneous sampling, camera trap distance sampling, and the association model, rely on similar types of data, but do not seem identical. In this article, I explore the relationships between different density estimators, including the random encounter model, the random encounter and staying time model, the time in front of camera approach, the time-to-event model, camera-trap distance sampling, the association model, and the space-to-event model. I show how these different estimators are related under two simplifying assumptions (perfect detectability, and animals moving as molecules in an ideal gas). I develop a map of mathematical relationships between these estimators. This framework helps readers understand how these methods are interconnected, providing a clearer conceptual foundation for selecting and implementing density estimation studies.

## Introduction

Historically, camera trap methods for estimating animal population size have primarily focused on marked animals, such as large carnivores with identifiable fur patterns (e.g. Karanth and Nichols, 1998). However, employing camera traps for abundance estimation remains a considerable challenge when dealing with unmarked populations. While N-mixture models can be used for this purpose, defining the effective sampling area of cameras is difficult (Gilbert et al., 2020). In fact, many methods have specifically been developed in the last two decades to estimate the size of a population of unmarked animals using camera traps.

Following a distinction established by Hutchinson and Waser (2007) in a general ecological framework and subsequently adapted by Campos-Candela et al. (2018) to camera-trap monitoring, I categorize density estimation methods into two main families: methods relying on animal-traps encounters and methods relying on animal-traps associations (Fig. 1). Animal-traps encounters can be defined as the full passage of an animal through the field of view of a camera trap. Each encounter is inherently defined in time, possessing a start, an end, a measurable duration, and an associated path length. These characteristics can be measured using traps equipped with movement sensors. Numerous population size estimators rely on the number of encounters detected by the traps. This is for example the case of the random encounter model, to my knowledge the first method proposed to estimate unmarked population size with camera trap data (Rowcliffe et al., 2008). The main challenge of encounter-based methods is that more active animals generate more encounters with traps, so that the unbiased estimation of population size requires the knowledge of the mean travel speed of animals – the so-called day range. Several methods have been proposed to estimate this day range using data collected on animal-traps encounters (e.g. Palencia et al., 2021, 2019). The alternative random encounter and staying time (REST) model uses the encounters duration measured by the traps to implicitly account for the mean travel speed of animals (Nakashima et al., 2017). Building on the same principle of using the total time spent in the field of view to avoid the need for speed estimation, other authors have developed the Time spent In Front of Cameras approach (Becker et al., 2022; Warbington and Boyce, 2020).

**Figure 1.**
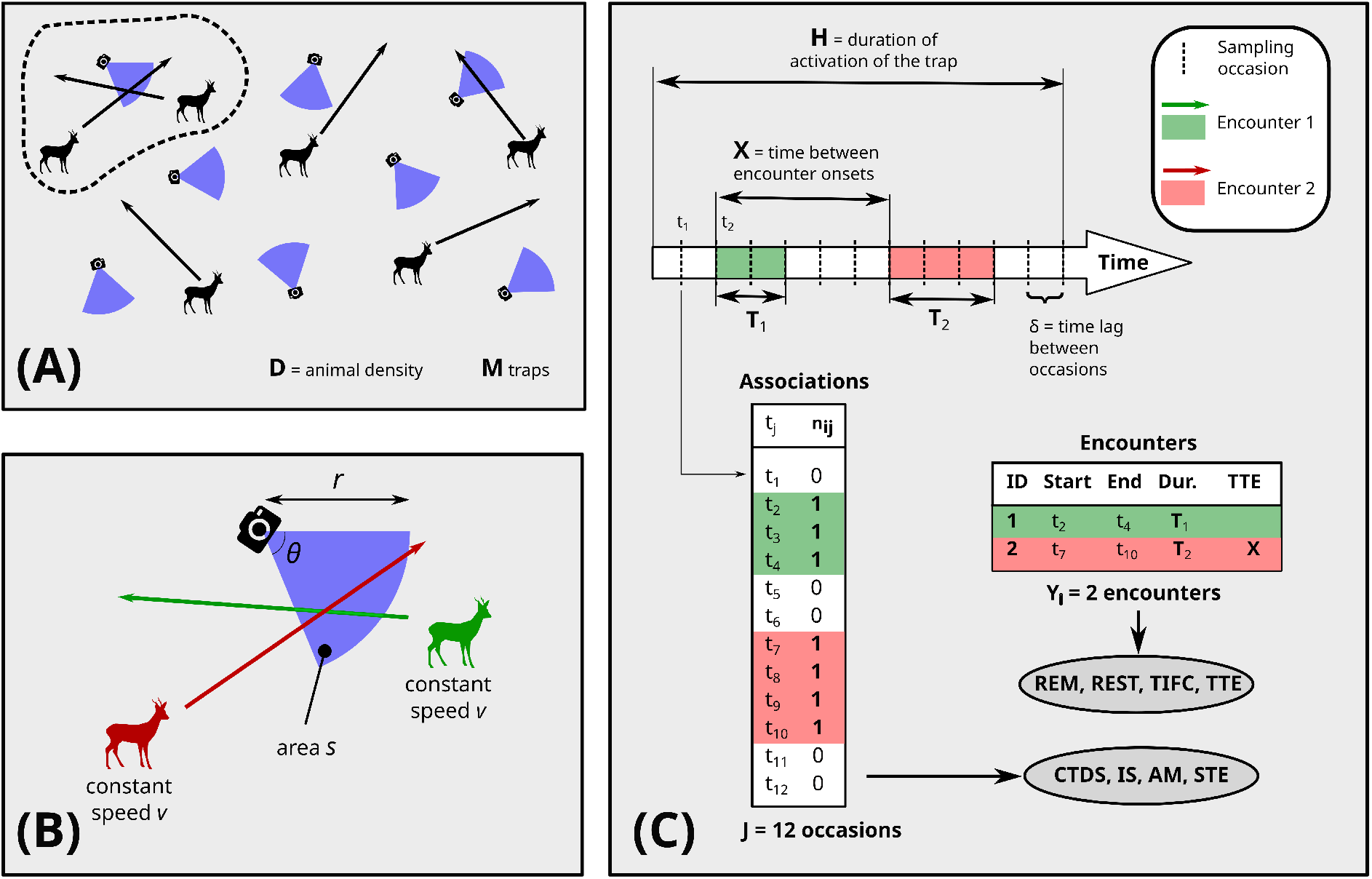
Schematic representation of a camera trap monitoring to estimate the animal density *D* in an area with *M* camera traps. (A) random distribution of the traps on the area and animals moving as molecules in an ideal gas. (B) Zoom on the upper left trap (circled with a dashed line in (A)): The trap with angle *θ* and radius *r* (which define its field of view) is crossed by two animals at two different times. (C) Data collection process for the trap described in (B): the camera trap is here set up to collect one photograph every *δ* seconds (i.e., sampling occasions). Using this setup, one can either record the number *n*_*ij*_ of animals in the field of view at every occasion (associations), or the encounter characteristics (start or onset, end, duration, time-to-event TTE = time elapsed between the two encounter onsets). Note: we suppose here that encounters start and end exactly at the time of sampling occasions for simplicity. This may not to be the case in other studies. The green and red encounters in (B) are displayed on the timeline and the two resulting datasets. Each dataset can be used to estimate density with a set of specific methods (REM = random encounter model, REST = random encounter and staying time, TIFC = time in front of camera, TTE = time-to-event model, CTDS = camera-trap distance sampling, IS/AM = instantaneous sampling/association model, STE = space-to-event model).

In contrast, methods relying on animal-trap associations do not require the explicit or implicit use of this day range to estimate population density. An animal-trap association is an instantaneous observation defined by the presence of an animal in the camera’s field of view at a specific time point. Conceptually, an association represents a point event in time, whereas an encounter is defined as the aggregation of successive associations, effectively transforming a series of instantaneous states into a measurable temporal duration. Associations typically occur when camera-traps operate in time-lapse mode, taking image at regular time intervals. The number of associations at a given time and place corresponds to the number of animals detected in a single image frame captured by a trap (Moeller et al., 2018). The total number of associations in the field of view of all camera traps at a given time is, on average, proportional to the population density. Several methods, such as the instantaneous sampling (Moeller et al., 2018), the association model (Campos-Candela et al., 2018), or the camera-trap distance sampling (Howe et al., 2017), rely on this relationship.

Moeller et al. (2018) proposed two additional methods that, while not directly relying on the number of associations or encounters, still leverage these concepts. The time-to-event approach relies on encounters: it uses the time elapsed between the start of a sampling period and the first encounter detection to estimate population density (the longer this duration, and the lower the density). The space-to-event approach, on the other hand, relies on associations detected using cameras operating in time-lapse. It estimates population size by sequentially sampling traps at a given time until an association is detected, using the cumulative area of the sampled traps to estimate population density (the larger this area, and the lower the density).

This plethora of estimation methods, while sharing a common foundation, are characterized by a large range of analytical requirements that can be confusing for newcomers. Although a single study design (e.g., a random grid of camera traps) is often compatible with several approaches, these methods rely on different components of the recorded data. Specifically, these differences manifest in: (i) the format of the input data required (e.g., encounters, associations, photographs or videos), (ii) the auxiliary information needed (e.g. the random encounter model requires the day range, while camera trap distance sampling does not), (iii) the factors accounted for (environmental variability, animal detectability).

Rather than focusing on the differing contexts in which these methods were developed, I aim in this paper to identify how they are related, and under which assumptions their estimators can be considered equivalent. Indeed, newcomers to this field (and even experienced biologists) often lack a global view of how these approaches are interconnected. Understanding the conceptual and practical links between their parameters would allow biologists to make more informed choices when selecting the most suitable approach for density estimation. For example, why do some methods require the average travel speed of animals and not others, despite relying on the same camera-trap data? Why does the time-to-event approach require the knowledge of the time required to cross the field of view of camera traps, but not the very similar space-to-event? What are the relationships between the different methods relying on associations? This comparative approach will help the reader to understand the theoretical connections between the different methods, and reveal potential avenues for combining different estimation techniques.

To establish a common ground for these various approaches, I chose to compare the different estimators under the ideal gas model. This model, originally developed by physicists to describe the collision rate among molecules in an ideal gas, has since been adapted by biologists to describe the movements of animals in a 2D space (Hutchinson and Waser, 2007). Although a version of this model exists where the speeds are drawn from a Maxwell-Boltzmann distribution, I consider here the version where animal speed is constant both over time (within individuals) and across all individuals in the population. This model further assumes that all individual paths are linear, with an initial uniform distribution of angles across individuals relative to a reference direction. While primarily developed to describe animal-animal encounter rates, this model has subsequently proved useful for describing the encounter rate between animals and camera traps (Rowcliffe et al., 2008). Ultimately, the simplicity of this model allows us to show that the density estimation methods – which use the same type data (camera traps) and share the same estimation target (animal density) – are fundamentally unified when considered within this framework.

In this paper, I present a map of the relationships between the different estimators discussed above (Fig. 2). My aim is to demonstrate the mathematical similarities between the different estimators under the assumptions of perfect detectability and of animals moving according to the ideal gas model used by Rowcliffe et al. (2008). This provides an intuitive understanding of how these methods are interconnected. To keep the paper as concise as possible, I have included the mathematical developments in the appendix.

**Figure 2.**
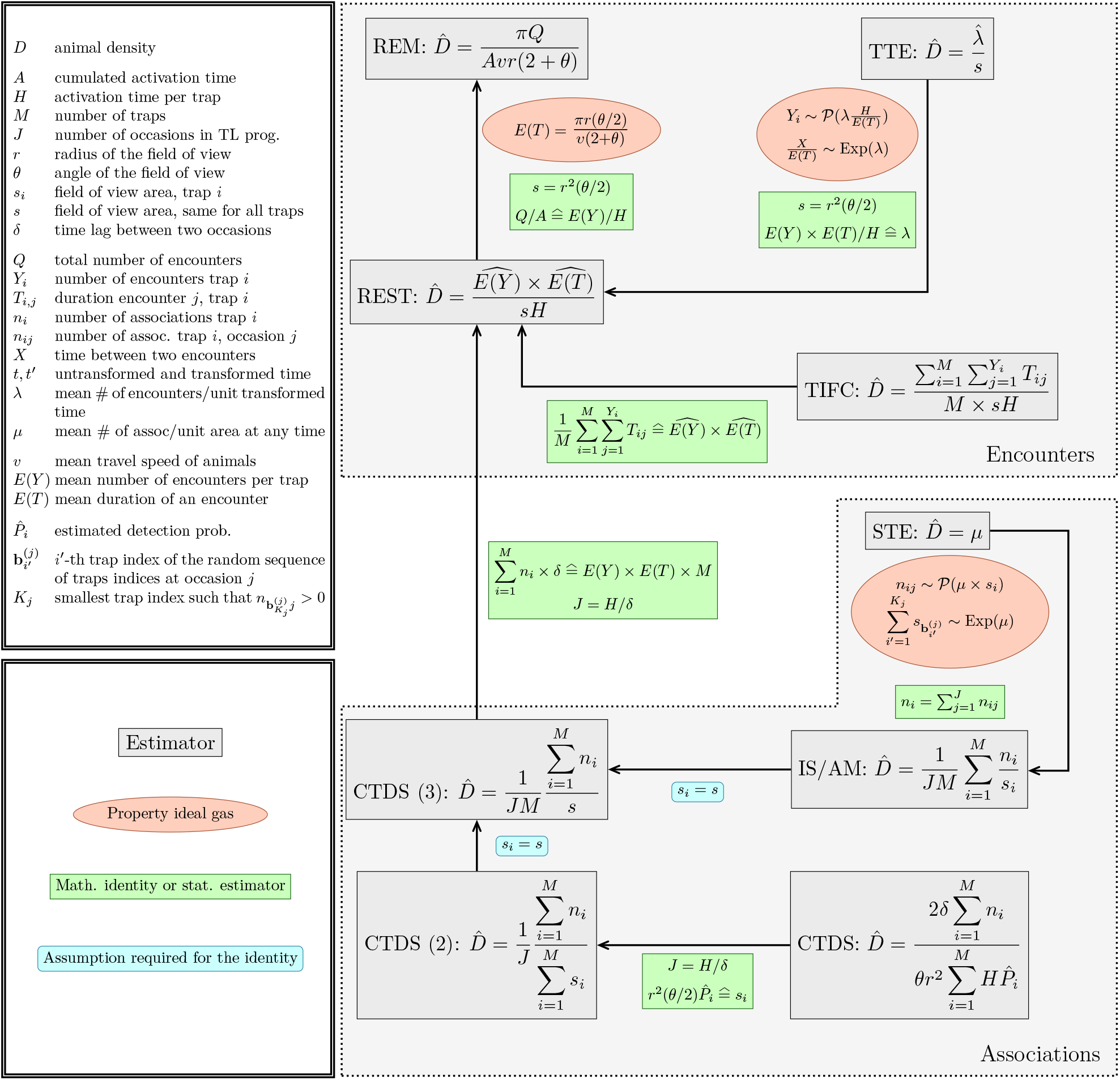
Map of the mathematical relationships between different density estimators based on the camera trap monitoring of a population of unmarked animals. Each grey square corresponds to an estimator (REM = random encounter model, REST = random encounter and staying time, TIFC = time in front of camera, TTE = time-to-event model, CTDS = camera-trap distance sampling, IS/AM = instantaneous sampling/association model, STE = space-to-event model). I distinguish association-based methods and encounter-based methods. For each pair of estimators, I indicate close to the arrow the conditions under which the two estimators are identical. I distinguish the properties of the ideal gas model, the mathematical identities/statistical estimators (≙ means that the right-hand side can be estimated by the left-hand side), and the assumptions required for the identity to hold. The inset at the top-right gives the meaning of the mathematical notations. The three estimators, CTDS, CTDS (2) and CTDS (3) correspond respectively to the original CTDS estimator, the CTDS estimator reformulated as a function of *J* and *s*_*i*_, and the CTDS estimator under the assumption of perfect detectability. The acronym TL stands for camera traps operating in time-lapse.

### Understanding the map

#### General methodology

In this paper, I study the relationships between the population size estimators corresponding to the 8 following methods: the random encounter model, the random encounter and staying time model, the time in front of camera approach, the time-to-event model, the space-to-event model, the instantaneous sampling approach, the association model, and camera-trap distance sampling.

One difficulty is that these estimators were developed in different statistical contexts, focusing on different aspects of the density estimation. Thus, some approaches rely on model-based estimators (e.g. random encounter and staying time, time-to-event, space-to-event, association model), while others, such as camera trap distance sampling, rely on design-based estimators.

Model-based estimators require distributional assumptions about the collected data, and often rely on maximum-likelihood estimation (e.g. some applications of the time-to-event approach suppose that the time between successive encounters follow a gamma distribution, Moeller et al., 2018). Conversely, design-based estimators require no distributional assumption on the data (Howe et al., 2017). Moreover, the different authors emphasise different factors that may bias the density estimation if unaccounted for. For instance, some estimators specifically account for imperfect detectability (e.g., camera trap distance sampling, Howe et al., 2017). Others account for the limited recording time predefined in most commercial camera traps, that prevents the precise recording of encounters duration (e.g., random encounter and staying time, Nakashima et al., 2017). Others, still, stress the importance of modelling the spatial heterogeneity in density (e.g., time-to-event, Moeller et al., 2018).

Rather than emphasising the differences, I would like to highlight how the different approaches, which all rely on camera trap data and share the goal of estimating animal density, are related to one another. I consider a simplified scenario of camera trap monitoring with perfect detectability (i.e., the camera trap is always triggered when the animal crosses its field of view). While this assumption is unrealistic in practice, it is useful to understand the connections between the estimators. I also suppose that all traps shared a uniform sampling period *H* (also called activation time below), meaning that they were active and ready to be triggered for the same duration. To facilitate their comparison, I reformulate the estimators as simple formulas rather than as the result of more complex estimation procedures such as the maximum likelihood. As noted previously, I also consider the simplistic case where animal movements are generated by the ideal gas model (Hutchinson and Waser, 2007), i.e. I assume that each individual moves at a speed that is constant both across individuals and over time, in a direction randomly drawn from a uniform circular distribution.

The map is presented in Fig. 2. The rest of the paper aims at giving the details that help to understand it.

#### The Random Encounter Model, Random Encounter and Staying Time, and Time In Front of Camera approaches

Rowcliffe et al. (2008) demonstrated how it is possible to use the ideal gas model to estimate the population density *D* from a function of the number of encounters *Q* (summed across all traps) during the total activation time *A* of the traps (also summed across all traps), the characteristics *r, θ* of the camera traps (resp. length and angle of their field of view), and the mean travel speed *v* of animals:

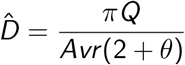

In the appendix, I show that under the ideal gas model, it is also possible to calculate the mean duration *E* (*T*) of an encounter (staying time) from the mean travel speed *v* of animals and the characteristics *r, θ* of the camera trap:

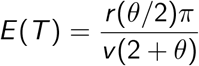

Note also that the overall encounter rate *Q/A* is also equal to:

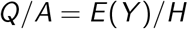

where *E* (*Y*) is the mean number of encounters per trap, and *H* is the duration of the study period (i.e., the activation time per trap). Finally note that the area of the trap field of view is *s* = *r* ^2^(*θ/*2) (Rowcliffe et al., 2008). Then, by expressing the density as a function of *E* (*T*) instead of *v*, and replacing *Q/A* with *E* (*Y*)*/H*, and simplifying, the density can simply be expressed as:

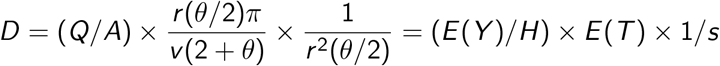

The random encounter and staying time estimator developed by Nakashima et al. (2017) is based on this last equation: it uses the data collected on the encounters by the traps to estimate *E* (*Y*) and *E* (*T*) and derive a density estimate, avoiding the need to estimate *v*. Note that the original model is actually more complex as it accounts for the possibility of right-censored staying time (addressing the limitations of commercial video traps, which stop filming after a predetermined time).

The conceptual basis of the random encounter and staying time model is also shared by the Time In Front of Camera approach (Becker et al., 2022; Warbington and Boyce, 2020). Indeed, the product *E* (*Y*) × *E* (*T*) corresponds to the mean time spent by animals in front of a camera per trap. If we consider the *M* traps of the study, this mean time can be estimated by the total duration of all encounters divided by the number of traps:

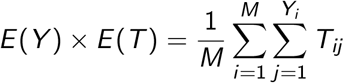

Where *T*_*ij*_ is the time spent by the animal in front of the trap *i* during encounter *j*. By substituting this into the previous equation, the density can be expressed as:

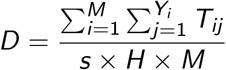

As noted by Warbington and Boyce (2020) and Becker et al. (2022), this highlights the close mathematical connection between the time in front of camera and the random encounter and staying time.

#### The Time To Event approach and its relationship with the Random Encounter and Staying Time approach

Moeller et al. (2018) developed a model-based approach to estimate the density based on the waiting times between arbitrary time points and the subsequent encounter onset (defined as the exact moment an animal enters the detection zone). The underlying principle is that shorter waiting times reflect higher animal densities. In their framework, the study period is divided into several sampling intervals, where the start of each interval serves as the reference point from which the waiting time is measured. Then, these authors discretize these intervals into sampling periods, and the waiting time is measured as a number of elapsed periods. However, as discussed in appendix, this discretization introduces small inconsistencies between the discrete time-to-event variable and the assumed continuous distribution. To ensure mathematical rigour within the scope of this study, I chose to maintain a continuous-time framework. While the full modelling approach by Moeller et al. (2018) accounts for numerous factors such as the heterogeneity of the density or imperfect detection, my objective here is not to provide a comprehensive alternative TTE estimator. Instead, I adopt a simplified version under the ideal gas model (assuming perfect detectability and homogeneous density) specifically to illustrate the relationships between the time-to-event and the random encounter and staying time frameworks.

First, like Moeller et al. (2018), I discretize the study period into sampling intervals (e.g. lasting one week). However, instead of discretizing each interval into sampling periods, I transform the time variable to *t*^*′*^ = *t/E* (*T*), maintaining its continuous nature. Note that the mean encounter duration *E* (*T*) is assumed to be known *a priori* (e.g., measured using the data collected from the camera traps). This transformation ensures that, on average, an encounter lasts exactly one unit of transformed time.

I demonstrate in the appendix that under the ideal gas model and with this transformation, the number of encounter onsets occurring with a camera trap during one unit of transformed time is Poisson distributed with mean equal to *λ* = *Ds*. In other words, the encounter onsets constitute a temporal Poisson process with parameter *λ*. A well-known characteristic of the Poisson process is that the time between two successive encounter onsets (inter-arrival time) – or equivalently, the time between the beginning of the sampling interval and the first encounter onset – follows an exponential distribution with a rate parameter also equal to *λ* and mean equal to 1*/λ* (Ross, 1996, p. 64). The time-to-event approach estimates *λ* from the observed time-to-event data by fitting an exponential distribution to these transformed times to the first encounter onset during each sampling interval. Then, exploiting the relationship between *λ* and the density *D* of animals under the assumed model, the time-to-event density estimator is:

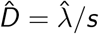

Now, *H* is the untransformed activation time of a trap. Based on the time transformation *t*^*′*^ = *t/E* (*T*), the transformed activation time of a trap is *H/E* (*T*). I established that the rate parameter *λ* = *Ds*. By definition, *λ* is also equal to the mean number of encounter onsets per unit of transformed time, which can be estimated by *E* (*Y*)*/*(*H/E* (*T*)). Replacing *λ* by this quantity in the time-to-event density estimator leads to the random encounter and staying time estimator (Fig. 2).

#### The Camera Trap Distance Sampling, Instantaneous Sampling, and Association Model approaches

The instantaneous sampling approach estimates the density using the number of animal-trap associations rather than the number of encounters (Moeller et al., 2018). The study period of length *H* is discretized into *J* instantaneous capture occasions, separated by a time lag of length *δ*. At each occasion *j*, the number *n*_*ij*_ of animals present in the field of view of the camera trap *i* is recorded. I define the number of associations per unit area detected by trap *i* at time *j* as the quantity *n*_*ij*_ */s*_*i*_, where *s*_*i*_ is the surface area of its field of view. An estimator of the population density is obtained by averaging the number of associations per unit area across all *M* traps and *J* occasions:

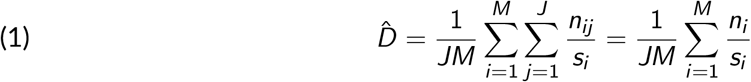

where 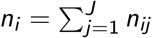.

Note that the association model of Campos-Candela et al. (2018), which estimates the density by averaging the number of animals counted per frame, is basically identical to the instantaneous sampling approach. The association model is actually a more complex modelling approach, accounting for environmental variations between cameras, but our assumption of the ideal gas model eliminates such variations, making the association model identical to the instantaneous sampling approach (see also equation (4) in Santini et al., 2022). I therefore refer to these two approaches as the instantaneous sampling/association model estimator.

The camera trap distance sampling approach is also similar to these estimators. Howe et al. (2017) showed how the distance sampling methodology, which accounts for imperfect animal detectability, can be used in the context of camera trap monitoring. These authors proposed first to measure the distance between detected animals and the camera traps for a random sample of associations, and subsequently to fit a detection function to these data by maximum likelihood. This statistical model describes how the number of associations decreases with increasing distance from the traps (e.g., see Buckland et al., 1993, p. 41 for a guide to select a suitable model form for this function). Under the assumption of random placement of camera traps, the animals density should not vary with distance to the trap. Thus, the observed change in detection density with distance reflects the reduction in animal detectability as the distance increases. From this detection function, an overall probability of detection 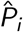 for the whole field of view can be estimated.

Then, the classic distance sampling estimator of the density, accounting for this imperfect detectability, can be used:

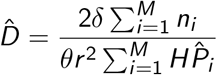

To get a better understanding of the relationship of this estimator with other estimators, I modify it slightly. Note that *s* = *r* ^2^*θ/*2 is the surface area of the field of view of the camera trap. Moreover, the effective surface *s*_*i*_ of the trap *i* can be estimated with 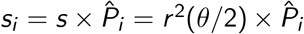. In addition, by definition, the number of occasions *J* = *H/δ*, so that the camera trap distance sampling estimator becomes:

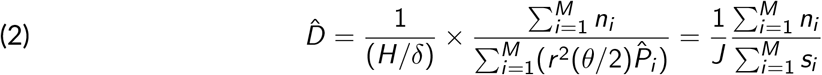

Comparing this equation with equation 1, we can see that the camera trap distance sampling and instantaneous sampling/association model estimators are not identical. The camera trap distance sampling estimator is equal to the ratio of the number of associations summed over all traps for each occasion divided by the sum over all traps of the effective surface of the traps. It is a ratio estimator (Thompson, 2002, p. 68). On the other hand, the instantaneous sampling/association model estimator is the mean across all traps of the ratios of the number of associations per trap divided by the surface of the trap. It is a mean-of-the-ratios estimator (Thompson, 2002, p. 84). In design-based estimation, it is well-known that these two estimators are biased. Although formulas exist to calculate the bias of both the ratio estimator (eq. 6.34 in Cochran, 1977, p. 161) and mean-of-the-ratios estimator (eq. 6.73 in Cochran, 1977, p. 174), it remains challenging to identify the optimal choice a priori. The optimal choice will depend on the existing correlation between the detectability and the animal density (i.e., the bias of both estimators increase when the density is larger in areas where animals are less detectable), the variability of detectability across camera traps, the form of the statistical distribution of the density, the number of camera traps, and the sampling design. In general, the ratio estimator is typically less biased than the mean-of-ratios estimator (as its bias decreases with the number of cameras, Cochran, 1977). Moreover, in locations where the detectability is low, the effective area *s*_*i*_ is small. This leads to a large variance of the ratio *n*_*i*_ */s*_*i*_ and, consequently, to instability of the mean-of-the-ratios estimator – a property not shared by the ratio estimator (Fewster et al., 2009). Actually, simulations may be required to decide which estimator is the most suitable in practice. Anyway, in our case, as I assume perfect detectability of animals, *s*_*i*_ = *s* and these two estimators become identical.

#### Relationship of camera trap distance sampling-related approaches with the random encounter and staying time approach

Palencia et al. (2021) demonstrated the equivalence of the random encounter and staying time estimator and the camera-trap distance sampling (see their appendix S1). I reproduce a similar rationale here to show how this equivalence – between instantaneous sampling/association model/camera trap distance sampling estimators and random encounter and staying time model under certain conditions – fits into our theoretical framework (Fig. 2). As before, I assume that all trap fields of view have equal surface areas. This simplifies the three association-based estimators to a function of the total number of associations detected during the study (estimator CTDS (3) on Fig. 2):

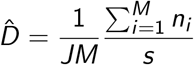

The main challenge in demonstrating the equivalence between the random encounter and staying time and camera trap distance sampling/instantaneous sampling/association model (CTDS / IS / AM) estimators arises from the difference in the data that they require: the random encounter and staying time estimator considers encounters, which have a duration, while the CTDS / IS / AM estimator focuses on instantaneous associations (point events in time). To bridge this conceptual gap, I assume that the time lag *δ* separating two successive occasions is very small (e.g., typically of a few seconds in camera trap distance sampling studies, see Howe et al., 2017). This small *δ* is crucial because it allows us to conceptually transform each instantaneous association into a short, measurable period of length *δ* during which an animal is present in the field of view of the camera. This step, in essence, lends temporal dimension to the association data, thereby making possible the comparision with the duration-based random encounter and staying time approach. Under this assumption, the total duration of presence of animals within the field of view of all camera traps can be estimated by 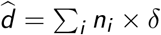.

Now, consider the design used for the random encounter and staying time approach, relying on animal-trap encounters. The mean time spent by animals in the field of view of a random camera trap is *E* (*Y*) × *E* (*T*). Therefore, the total duration of presence of animals within the field of view of all *M* traps is equal to *d* = *M* × *E* (*Y*) × *E* (*T*). When demonstrating this equivalence, Palencia et al. (2021) illustrated this equality by giving the example of “a single three-second encounter in the [random encounter and staying time] context (Y = 1, T = 3) would be discretised at one second intervals in the [camera trap distance sampling] context as n = 3, *δ* = 1)”. Thus, we can substitute ∑_*i*_ *n*_*i*_ ×*δ/M* for *E* (*Y*) ×*E* (*T*) in the random encounter and staying time estimator, simplify, and arrive at the camera trap distance sampling estimator. This shows their equivalence (Fig. 2).

The fact that we can estimate *M* × *E* (*Y*) × *E* (*T*) by ∑_*i*_ *n*_*i*_ × *δ* clarifies why encounter-based methods require knowledge of the mean travel speed, while association-based methods do not. Faster moving animals results in more frequent but shorter encounters, yet the total time spent in front of the traps remains constant.

#### The Space To Event approach and its relationship with the Instantaneous Sampling/Association Model approach

The Space-to-Event approach developed by Moeller et al. (2018) is conceptually similar to the time-to-event approach, but focuses on the space separating animals at a given time. The underlying idea is that a small density corresponds to large average amount of space between animals. Like the instantaneous sampling/association model and camera trap distance sampling approaches, the space-to-event model defines instantaneous sampling occasions separated by a given time-lag, *δ*. This model-based approach consists in defining a random sequence of the *M* installed camera traps for a given sampling occasion. The cumulative space without an association covered by the traps in the sequence, up until the first detection, is termed the space-to-event. This cumulative area is assumed to follow an exponential distribution with rate *µ*. This parameter corresponds to the mean number of animals per surface area unit, and is therefore a density estimator:

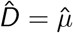

As for other approaches, I redevelop the space-to-event approach in the context of the ideal gas model. In an ideal gas, independent molecules are randomly distributed in space at any given moment (Hutchinson and Waser, 2007). This complete spatial randomness of independent events at instantaneous moments, observed at a fixed instant in time, characterizes the Poisson point process (Diggle, 1983, p. 50). In other words, by sampling at any time *j* an area *s*_*i*_, the resulting number of associations in this area follows a Poisson distribution *n*_*ij*_ ∼ 𝒫 (*µ*_*i*_), with *µ*_*i*_ = *D* × *s*_*i*_.

It is well-known that the squared distance between points (squared inter-event distance) generated by a Poisson point process follows an exponential distribution with parameter *π* × *D* (Diggle, 2013, p. 24). Or, equivalently, the squared distance *d* ^2^ between any trap on the study area and the closest animal is exponentially distributed with parameter *π* ×*D*. In other words, the largest amount of surface area *πd* ^2^ centred on the trap without any animal in it is exponentially distributed with parameter *µ* = *D*. This property forms the basis of the space-to-event approach.

Consider a given sampling occasion *j*. We form a sequence of trap indices *i* in a random order. The probability that the first trap in the sequence with an association (i.e., *n*_*ij*_ *>* 0) occurs at the *k*-th sampled trap is equal to the probability that the cumulated sampled space without an association is 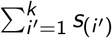 Here, *s*_(i ′)_ corresponds to the surface area of the field of view of the *i* ^*′*^-th trap of the sequence. This cumulative area is exponentially distributed with a rate parameter equal to the density *D*. Note that this approach presents the same small inconsistency as the original time-to-event approach of Moeller et al. (2018). Specifically, the measured space-to-event variable is essentially discrete, yet it is typically modelled using a continuous distribution. For the purpose of this study, I will follow Moeller et al. (2018) and ignore the discrete nature of this variable.

We can then form random sequences of traps for all sampling occasions, and measure the space-to-event for each occasion. We then fit an exponential distribution to the distribution of the amounts of area sampled at each occasion required before an association is detected. The estimated parameter of this exponential distribution is the estimated density. Note that the modelling approach of Moeller et al. (2018) actually accounts for the possibility of censored values for this area (i.e., the fact that there may be no associations at all with any trap at a given occasion).

Now, the relationship with the instantaneous sampling approach becomes clear: since at any time *j*, the number *n*_*ij*_ of associations with trap *i* follows a Poisson distribution with mean *Ds*_*i*_ under the ideal gas model, it follows that the ratio *n*_*ij*_ */s*_*i*_ – the number of associations per unit area detected by trap *i* at sampling occasion *j* previously – is a direct estimator of the true density *D*. The instantaneous sampling estimator, as previously derived, is obtained by simply averaging this number of associations per unit area across all sampling occasions and all traps (Fig. 2).

## Discussion

When I started working on camera trap data with my team, I was overwhelmed by the apparent multitude of potential approaches to estimate animal density. While these methods are fundamentally interconnected (and a single field protocol can often provide the data needed for several of them) the literature has frequently stressed their unique characteristics in terms of detection rates, spatial heterogeneity in density, available side information, etc. (see for example Gilbert et al., 2020; Santini et al., 2022). However, I missed a global view of how these approaches are interconnected, and why some some methods require certain data type (e.g. the average travel speed) that others did not. Building this map (Fig. 2) helped me to grasp the theoretical connections between different density estimation methods.

Note that this map is not exhaustive. Other approaches have been developed to estimate density, like the unmarked spatial capture recapture of Chandler and Royle (2013), which uses the spatial correlation between neighbouring traps to identify the activity centers of the home range of non-identifiable animals, and thereby derive a density estimate (although notoriously imprecise, see Gilbert et al., 2020; Wearn et al., 2022). I focused here on methods relying on the number of associations/encounters to estimate the density, because they are among the most widely used in the literature (e.g., Nakashima, 2022; Santini et al., 2022).

To allow this study of the relationships between estimators, I simplified some of them. For example, the random encounter and staying time estimator that I presented does not correspond exactly to the one presented by Nakashima et al. (2017). However, the random encounter and staying time estimator on Fig. 2 captures the spirit of the method, and helps to clarify how association-based methods and encounter-based methods are related.

To build this map, I further made two key simplifying assumptions: (i) animals move as molecules in an ideal gas model, and (ii) they are detected as soon as they enter the camera’s field of view (perfect detectability). These simplifying assumptions serve a purely didactic purpose: they allow us to clarify the intrinsic relationships between the parameters of the different methods. For instance, the ideal gas model assumption helps in understanding how animal speed and camera traps parameters (radius, angle) relate to the expected time in the field of view, how density connects with the time between encounter onsets, and how encounters translate into associations, justifying the use of a Poisson distribution for the spatial distribution of animals and temporal distribution of encounters. Similarly, although nearly all estimation methods can accommodate for imperfect detectability – either by replacing the surface area of the trap’s field of view with an effective area (e.g., the random encounter model), by integrating imperfect detection directly into the density estimation (e.g., the camera trap distance sampling), or by making assumptions on the effect of this imperfect detection on the distribution of modelled variable (e.g., the time-to-event approach) – the demonstrated equivalence between instantaneous sampling/association model/camera trap distance sampling/random encounter and staying time model is valid only under the assumption of perfect detectability. While this assumption is useful for theoretical unification, empirical applications must account for imperfect detection by accurately estimating the detection area. This remains a critical step, often requiring dedicated field tests with realistic targets to calibrate specific camera models under various conditions (see e.g., Apps and McNutt, 2018; Randler and Kalb, 2018).

I stress that the ideal gas assumption was only made for the purpose of this theoretical comparison; this assumption is not necessarily required by the methods themselves. Indeed, camera trap distance sampling, instantaneous sampling and association model do not rely on the ideal gas model assumption at all. Thus, Campos-Candela et al. (2018) demonstrated that, under certain conditions, the association model remains valid for animals exhibiting home-range behavior rather than purely random, unbounded movement. The random encounter and staying time is a model-based approach; to develop this model, Nakashima et al. (2017) made assumptions for the distribution of the time required to cross the field of view that do not imply the ideal gas model (though they still made other assumptions). The random encounter model is actually the only approach explicitly developed based on the ideal gas model assumption (Rowcliffe et al., 2008). However, this underlying model was only used for mathematical convenience, and the random encounter model approach can be shown to be robust to the violation of this assumption (Rowcliffe et al., 2013). Thus, our forthcoming work, based on simulations of a population of roe deer, shows that the random encounter model allows for unbiased population size estimates, despite individual variation in space use, non uniformly distributed home ranges, considerable habitat selection, and variable daily activity (Calenge et al., 2025).

However, our assumption of the ideal gas model made it easier to study the connections between the different methods. If this assumption no longer holds, some equivalences no longer hold. For example, we used this assumption to demonstrate the equivalence between the random encounter model and the random encounter and staying time model, and this equivalence is much harder to identify without it. Furthermore, the equivalence between the time-to event approach and the random encounter and staying time approach relies heavily on the theoretical connections between the Poisson process describing encounters (which depends on animal density) and the exponential distribution describing the waiting time before an encounter. This Poisson process is a direct result of the ideal gas hypothesis. The violation of this model would necessarily lead to a different distribution for the waiting time. If the ideal model no longer holds, there is no reason to believe that time-to-event and random encounter and staying time approaches would lead to similar estimates. The same doubts can be raised for the space-to-event approach, which strongly relies on the hypothesis that the animals locations are distributed as a spatial Poisson point process at any time. Although some studies suggested that the time-to-event approach is robust to departure from this model (Loonam et al., 2021), other work, such as Gurarie and Ovaskainen (2011), shows that the time-to-event (that they call “first hitting time” with a random target) is smaller when movements are more tortuous, suggesting that the equivalence with other encounter-based method might not hold under different movement models.

Note that even if strong theoretical connections exist between these estimators, and even if two of them are similar under the ideal gas model, this does not imply practical equivalence in real-world studies. In particular, animal behavior may affect the precision of certain estimates more than others. For instance, animals activity cycles and variation in movement speed will lead to increased variance in encounter rates. This variability may impact the precision of the different estimators differently, although further studies are required to precisely quantify this effect. Similarly, the presence of habitat selection and animal interactions, resulting in aggregated populations can strongly affect the efficiency and, in particular, the precision of the estimation methods (Gurarie and Ovaskainen, 2013; Howe et al., 2017; Palencia et al., 2022), with the magnitude of this effect likely varying between approaches.

The present work did not consider the different methods available to estimate the variance of the density estimation (e.g., bootstrap, asymptotic properties of maximum likelihood, design-based estimator). Given the multitude of available variance estimation methods, both the practical aspects of their implementation (e.g., bootstrap is often easy to implement) and their performance must be considered. More generally, regarding precision, simulations of camera-trap monitoring of a population with simulated animal movements can help to assess the relevance of camera-trap monitoring and to determine the most suitable density estimator under specific conditions (Nakashima, 2022; Santini et al., 2022).

An important implication of the ideal gas model is that, by definition, any placement of the camera traps on the study area leads to random movement of animals with respect to the cameras. For this reason, we ignored the sampling design in building our map, and implicitly assumed a random placement. However, as soon as the movements are no longer completely random across the area, the issue of the optimal sampling design arises. A crucial characteristic of all estimation methods considered in this paper is their assumption that animal movements are random with respect to the cameras. For this reason, most of them assume a random placement of the camera traps across the study area, possibly using a stratified sample to account for the variable density resulting from habitat selection and aggregated animal distribution. This requirement implies placing camera traps even in poor habitat, which many biologists may perceive as a waste of effort. Many prefer to place camera traps only in good habitat to maximize the detections. However, this preference would likely result in an overestimation of the density, and may even bias the estimation of population trends (Calenge et al., 2025). The sampling design issue is a complex matter that must be taken seriously when designing a camera trap monitoring.

Although I focused on the similarities between methods, the user will, of course, also need to consider their differences when choosing the appropriate approach. This choice depends on their specific goals and requirements. While several studies provide valuable guidance (Gilbert et al., 2020; Nakashima, 2022; Santini et al., 2022), the map in Fig. 2 offers a complementary perspective. For example, one common criticism with the association model is that it does not allow to account for imperfect detectability (Santini et al., 2022). However, the map shows here that this approach is, in its principle, very similar to the instantaneous sampling and camera trap distance sampling, which account for this imperfect detectability. So that if the user is interested in association-based methods accounting for detectability, they can consider instantaneous sampling and/or camera trap distance sampling. The choice between instantaneous sampling and camera trap distance sampling depends on whether the user prefers a ratio estimator (camera trap distance sampling) or a mean-of-ratio estimator (instantaneous sampling), and simulations can be used to assess the bias of each method in a specific context.

The choice of the best method also depends on the elements available to the user (Gilbert et al., 2020; Wearn et al., 2022): whether data on the encounters duration are available or not, whether encounter or association data are collected, whether encounter data can be transformed into association data, etc. Thus, the choice between instantaneous sampling and camera trap distance sampling also depends on whether distance data are available to fit a detection function (camera trap distance sampling) or not (instantaneous sampling). Note however that while the distance sampling estimator is typically used with detection probabilities estimated from detection functions fitted to distance data, it can be adapted to accommodate detection probabilities derived from other methods (e.g., from field experiments).

In summary, the optimal estimation method depends on a complex interplay of factors: the size of the study area, the financial and human resources determining the number of traps and the feasibility of experiments to estimate detectability at different trap positions, animal density and behaviour (home-range size, habitat selection, social interactions, activity cycle), the correlation between detectability and animal density, and the possible sampling design. I do not believe that the map identifies which is the best estimator, as I do think that there is no one-size-fits-all solution. Consequently, this work does not aim at completely solving the issue of the optimal method to be used in every case. However, it offers a rigorous framework for biologists interested in estimating animal densities, as well as practitioners responsible for designing monitoring programs. By revealing how these different frameworks are theoretically interconnected, this study provides more than a conceptual map; it offers a clearer view of the methodological landscape and a deeper intuitive understanding of the core concepts. Ultimately, this unification can meaningfully inform study design and estimation by fostering a heightened awareness of the mechanisms and assumptions that underpin each of these approaches.

## Acknowledgements

I would like to warmly thank Mathieu Garel, Sonia Saïd, Jules Chiffard and Maryline Pellerin, as well as my colleagues of the MAS network of the French biodiversity management organisation (*Office français de la biodiversité*) for the fruitful discussions and advices regarding the estimation of animal density with camera trap data. I am also grateful to Simon Lacombe, Elie Gurarie, Frédéric Barraquand and an anonymous referee for providing very constructive and useful comments on previous versions of this manuscript, which significantly helped to improve its quality.

## Fundings

The author declare that he has received no specific funding for this study.

## Conflict of interest disclosure

The author declare that he complies with the PCI rule of having no financial conflicts of interest in relation to the content of the article.

## Data, script, code, and supplementary information availability

Supplementary information is available online, on the same bioRxiv page as the preprint (https://doi.org/10.1101/2025.02.28.640755).

## Notes

### Competing Interest Statement

The authors have declared no competing interest.

### Summary of Updates

This version is identical to the previous one, but I now added a badge on the first page indicating that it has been recommended by PCI Ecology (https://doi.org/10.24072/pci.ecology.100791).

