## Appendix for "Camera trap monitoring of unmarked animals: a map of the relationships between population size estimators"

Clément Calenge

In this appendix, I show how the ideal gas model can be used to estimate the mean staying time $E(T)$, i.e., the mean duration of an animal-trap encounter, as well as the mean path length $\bar{\ell}$ during an encounter. Then, I show that the number of encounters occurring during time $t$ is a Poisson process with encounter rate equal to $Ds$. Finally, I discuss the difficulties introduced by the time discretization in the development of the time-to-event approach by Moeller et al. (2018).

#### 1 Mean staying time and mean encounter length

I use the same mathematical notations as in the paper. Figure 1 summarize the rationale. We derive expressions for these two quantities more formally below.

First, note that the probability of an animal entering a camera trap's field of view is the same, regardless of whether the animal is moving toward the trap or the trap is moving toward the animal at speed $v$. Consider the case where the trap is moving with speed $v$ at an angle $\alpha$ relative to the North direction. We define the *profile* of the trap as the maximum width $L_\alpha$ of the field of view perpendicular to the direction of movement (Fig. 1C). Since the profile is uniquely defined by the angle $\alpha$ of the velocity vector to which it is perpendicular, we can identify a profile just by using this angle. For example, we can consider a given profile $\alpha$, corresponding to the maximum width $L_\alpha$.

The path length in the field of view of the trap depends on the entry point of the animal. We define a Cartesian coordinate system with the abscissa axis parallel to the trap's profile (Fig. 1D). Let $l(x;\alpha)$ be the length of the animal path entering the field of view at the abscissa coordinate $x$ defined by the profile $\alpha$ of the trap.

The surface area $s$ of the field of view of the trap is equal to the integral over $x$ of this length:

$$s = \int l(x;\alpha)dx$$

Since the profile width is equal to $L_\alpha$, then it follows that

$$\bar{l}_\alpha = \frac{1}{L_\alpha}\int l(x;\alpha)dx = s/L_\alpha$$

corresponds to the mean length of the path of the animal that would enter the field of view with a velocity vector of orientation $\alpha$ (this mean is calculated for all possible entry points).

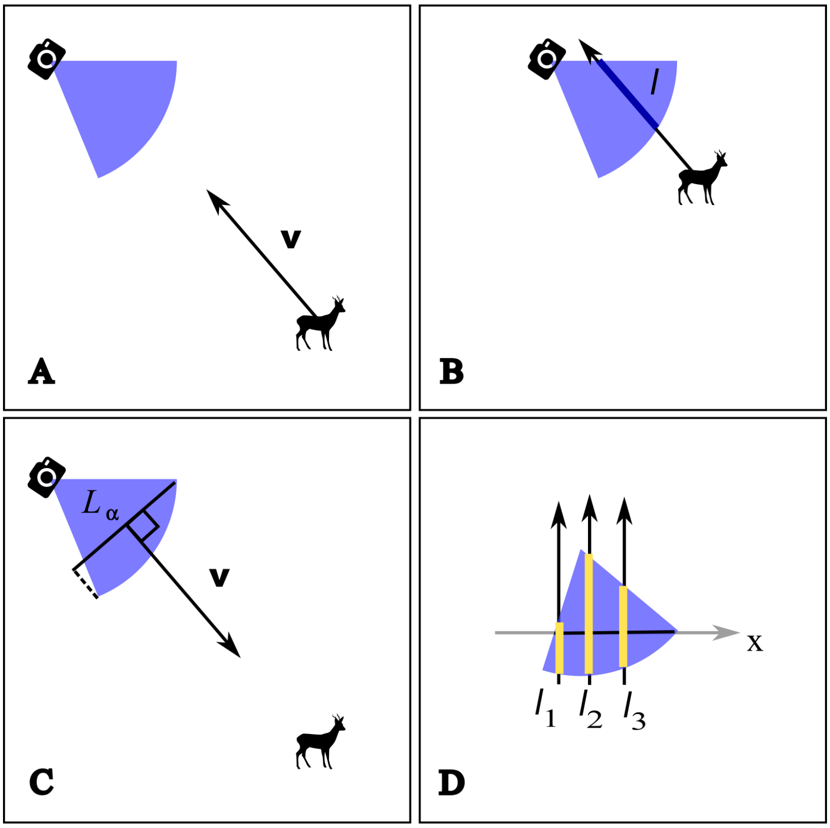


Figure 1: Calculation of $E(T)$ and $\bar{\ell}$ with the ideal gas model: (A) consider the case of an animal moving at constant speed $v$ and constant orientation toward the trap . (B) The staying time $T$ can be calculated as $l/v$, with $l$ the path length within the field of view of the trap. (C) From a mathematical point of view, it is equivalent to consider whether the animal or the trap is moving (relative velocity is the same in the two cases). Consider the case where the trap is moving, and note $L_\alpha$ the maximum width of the profile orthogonal to the velocity vector. We define the angle $\alpha$ characterizing $L_\alpha$ as the angle between the velocity vector $\mathbf{v}$ and the north direction. (D) The length $l$ of the path traversed by the animal in the field of view of the trap depends on its entry point. However, for a given velocity vector, the different possible paths corresponding to the different entry points are parallel, i.e., perpendicular to the profile of the trap. We define a new coordinate $x$ along the this profile of maximum width. For all entry points with a coordinate $x_i$ in the field of view, there is a path length $l(x_i;\alpha)$ (on this figure, this notation is simplified to $l_i$). Note that $\bar{l}_\alpha = \frac{1}{L_\alpha}\int l(x;\alpha)dx$ corresponds to the mean path length of the animal. The integral over $x$ of their length, $s = \int l(x;\alpha)dx = L_\alpha \times \bar{l}_\alpha$, corresponds to the area of the field. Note that this area can also be calculated from its length and angle. Therefore, one can also calculate the mean path length of the animal in the field with this profile by $\bar{l}_\alpha = s/L_\alpha$. If $v$ is the trap speed, the mean duration is $T_\alpha = \bar{l}_\alpha/v$, that is, $T_\alpha = s/(v \times L_\alpha)$. By averaging $T_\alpha$ over all possible $\alpha$, it is possible to estimate $E(T)$. The mean path length $\bar{\ell}$ can then be derived by $v \times E(T)$.

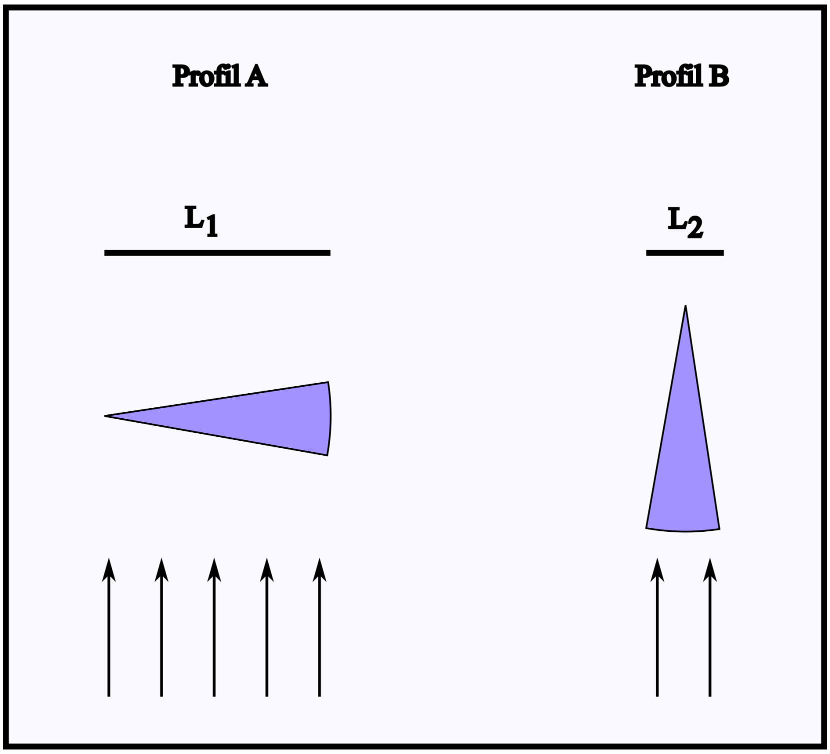


Figure 2: Illustration of the need to weight the mean path lengths in the field of view of the trap by the profile width. In the case (A), the field of view is characterized by a profile with maximum width $L_1$. In the case (B), the field is characterized by the narrowest possible profile (width $L_2$). The animal has more chances to traverse the field of view in (A) than in (B). When calculating the mean path length over all profiles, we need to give more weight to the mean length associated to profile A that to the mean length associated to profile B. We therefore need to weight these averages by the profile widths.

Note that the surface area of the field of view is equal to:

$$s = \pi \times r^2 \times \theta/(2\pi) = r^2 \times \theta/2 \tag{1}$$

with $\theta, r$ the angle and length of the field of view.

Given that the constant speed is equal to $v$, the mean duration $T_\alpha$ of the encounter corresponding to profile $\alpha$ is by definition equal to $\bar{l}_\alpha/v$, that is:

$$T_\alpha = \frac{s}{v \times L_\alpha}$$

The expectation of $T_\alpha$ over all possible profiles can be calculated by integrating $T_\alpha$ over all possible values of $\alpha$, weighting each value of $T_\alpha$ with $L_\alpha$ to account for the fact that there are more chances for an animal to cross the field of view orthogonally to a large profile than orthogonally to a narrow profile (Fig. 2).

In other words, in the calculation of the values of $T_\alpha$, we weight each value of $T_\alpha$ with:

$$\frac{L_\alpha}{\int_0^{2\pi} L_{\alpha'} d\alpha'}$$

The expectation of the encounter duration (mean staying time) is therefore:

$$\begin{aligned} E(T) &= \int_0^{2\pi} \frac{L_\alpha}{\int L_{\alpha'} d\alpha'} \times \frac{s}{v \times L_\alpha} d\alpha = \frac{s}{v \times \int L_{\alpha'} d\alpha'} \int_0^{2\pi} d\alpha \\ &= \frac{s \times 2\pi}{v \times \int L_{\alpha'} d\alpha'} \\ &= \frac{s}{v \times \left( \frac{1}{2\pi} \int L_{\alpha'} d\alpha' \right)} \end{aligned} \tag{2}$$

Note that

$$\frac{1}{2\pi} \int_0^{2\pi} L_{\alpha'} d\alpha' = \bar{L}$$

corresponds to the mean maximum width of the profile calculated over all possible velocity vectors. Rowcliffe et al. (2008) have showed that this mean maximum width $\bar{L}$ can be calculated with:

$$\bar{L} = r \times (2 + \theta)/\pi$$

Replacing this equation in equation (2):

$$E(T) = \frac{s \times \pi}{v \times r \times (2 + \theta)}$$

And replacing the equation (1) giving the value of $s$ in this equation, we have:

$$E(T) = \frac{r^2 \times \theta/2 \times \pi}{v \times r \times (2 + \theta)} = \frac{r \times \theta/2 \times \pi}{v \times (2 + \theta)} \tag{3}$$

which corresponds to the expression used in the paper for $E(T)$.

Note that it is also possible to use these developments to give an expression for $\bar{l}$, the mean paths length per encounter:

$$\bar{l} = v \times E(T) = \frac{r \times \theta/2 \times \pi}{(2 + \theta)} \tag{4}$$

which corresponds to the expression used in the paper for $\bar{\ell}$.

#### 2 The encounters constitute a Poisson process

As in the previous section, consider the situation where an animal is stationnary and a camera trap is moving. Rowcliffe et al. (2008) demonstrated clearly that the average profile of a camera trap across all possible angles of approach is equal to:

$$r \frac{2 + \theta}{\pi}$$

Therefore, during time $t$ and given a movement speed equal to $v$, this average profile of this camera trap sweeps a surface equal to:

$$vtr\frac{2+\theta}{\pi}$$

Given a study area covering a surface equal to $S$, the probability that an animal sampled in the population falls in the area swept by this profile is:

$$\zeta = vtr\frac{2+\theta}{\pi S} \tag{5}$$

Thus, the occurrence of an encounter between a given animal and a given camera trap is a Bernoulli experiment with probability $\zeta$.

Now consider $N$ independent animals on the study area moving as molecules in an ideal gaz model, and a unique camera trap. As stated, the occurrence of an encounter between a specific animal and the camera trap is a Bernoulli experiment, and since the animals are independent, it follows that the number of encounters occurring during time $t$ is binomial $\mathcal{B}(N, \zeta)$. Since the probability $\zeta$ is typically very small (the area covered by a camera trap is small in comparison to the study area), and for large $N$, the number of encounters $Y_i$ with the trap $i$ converges towards a Poisson distribution:

$$Y_i \sim \mathcal{P}(N\zeta)$$

Now, since the density is equal to $D$, in average, the mean number of animals on the study area of surface $S$ is $N = DS$, so that:

$$Y_i \sim \mathcal{P}(DS\zeta)$$

Replacing $\zeta$ with equation 5, the expectation of this Poisson distribution is:

$$Dvtr\frac{2+\theta}{\pi}$$

We now make a change of variable, by transforming the time, as in the paper. We set $t' = t/E(T)$. With this change, one unit of transformed time corresponds to the average time required by the animal to cross the camera trap's field of view. Therefore, during one unit of transformed time, the expectation of the Poisson distribution is:

$$\lambda = DvE(T)r\frac{2+\theta}{\pi}$$

Replacing $E(T)$ with equation 3, we have:

$$\begin{aligned}\lambda &= Dvr\frac{\pi r(\theta/2)}{v(2+\theta)} \times \frac{2+\theta}{\pi} \\ &= Dr^2(\theta/2) \\ &= Ds\end{aligned}$$

With again $s$ the surface of the camera trap's field of view. Therefore, under the ideal gas model, the number of encounters $Y_i$ occurring with a camera trap during one unit of transformed time is Poisson distributed with mean $Ds$.

Since the counting process $N(t')$ giving the number of encounters occurring during time $t'$ is characterized by $N(0) = 0$, that increments are independent and that the number of encounters in any interval of length $t'$ is Poisson distributed with mean $\lambda t$, this counting process is a Poisson process (Ross , 1996, Definition 2.1.1, p. 59).

### 3 Difficulties introduced by the time discretization in the development of the time-to-event approach by Moeller et al. (2018)

In this section, we focus on punctual events representing encounter onsets (i.e., the exact moments an animal enters the detection zone). As demonstrated in the previous section, under the ideal gas model, these events constitute a homogeneous temporal Poisson process with rate $\lambda = Ds$.

The time-to-event approach of Moeller et al. (2018) relies on the principle that a high density of animals in a given area is associated with a short waiting time between an arbitrary time point and the first encounter onset (defined as the exact moment when an animal enters the detection zone after that time point). More precisely, the study period is divided into sampling intervals, the beginnings of which correspond to these arbitrary time points. Then, each sampling interval is divided into discrete sampling periods, each with a duration of $E(T)$. Note that $E(T)$ is assumed to be known a priori (e.g., measured using the data collected from the camera traps).

By discretizing time, the process describing the occurrence of encounters onsets is no longer a Poisson process, but a discrete-time stochastic process called a Bernoulli process (Gallager , 1996, p. 14). Since the process in continuous time is a Poisson process with rate $\lambda$ and mean $1/\lambda$, the probability $p$ that a discrete sampling period of duration $E(T)$ contains at least one encounter onset is:

$$p = 1 - \exp(-\lambda \times E(T))$$

That is, one minus the probability of having no encounter onset during a time interval of length $E(T)$. Consequently, within each sampling occasion, the probability distribution describing the time required to observe an encounter onset is a geometric distribution (Gallager , 1996, p. 15). The probability that the first encounter onset occurs at the j-th period is therefore:

$$p_X(j) = (1 - p)^{j-1} p$$

Therefore, when discretizing time, this approach should normally be used to describe the encounter process.

Note that in a Bernoulli process, as the duration of the sampling periods tends toward 0 and the number of sampling periods becomes large, the Bernoulli process converges toward a Poisson process, and the geometric distribution of encounter onsets converges toward an exponential distribution (Gallager , 1996, p. 69).

Moeller et al. (2018) do not account for this discretization in their model, and directly assume an exponential distribution for the time-to-event measured in discretized time. This creates a small inconsistency between the sampling design and the chosen distribution (note that the same type of inconsistency is observed for the space-to-event model).

As a follow-up study, it would be valuable either to explore the effect of this discretization on density estimates (and their precision), to adapt the approach of Moeller et al. (2018) by using a geometric distribution instead of an exponential one, or to adopt a framework similar to the one developed in my article, using continuous transformed time to measure the time to event.
